# An advanced automated patch clamp protocol design to investigate drug – ion channel binding dynamics

**DOI:** 10.1101/2021.07.05.451189

**Authors:** Peter Lukacs, Krisztina Pesti, Mátyás C. Földi, Katalin Zboray, Adam V. Toth, Gábor Papp, Arpad Mike

## Abstract

Standard high throughput screening projects using automated patch-clamp instruments often fail to grasp essential details of the mechanism of action, such as binding/unbinding dynamics and modulation of gating. In this study, we aim to demonstrate that depth of analysis can be combined with acceptable throughput on such instruments. Using the microfluidics-based automated patch clamp, IonFlux Mercury, we developed a method for a rapid assessment of the mechanism of action of sodium channel inhibitors, including their state-dependent association and dissociation kinetics. The method is based on a complex voltage protocol, which is repeated at 1 Hz. Using this time resolution we could monitor the onset and offset of both channel block and modulation of gating upon drug perfusion and washout. Our results show that the onset and the offset of drug effects are complex processes, involving several steps, which may occur on different time scales. We could identify distinct sub-processes on the millisecond time scale, as well as on the second time scale. Automated analysis of the results allows collection of detailed information regarding the mechanism of action of individual compounds, which may help the assessment of therapeutic potential for hyperexcitability-related disorders, such as epilepsies, pain syndromes, neuromuscular disorders, or neurodegenerative diseases.

## Introduction

Most small-molecule sodium channel inhibitors bind to the local anesthetic binding site, and they are strongly state-dependent, showing ∼10-fold to 1000-fold higher affinity to inactivated channels^1^. For this reason, as it has long been recognized, determining an IC_50_ value with a single voltage protocol means practically nothing. Radically different IC_50_ values can be measured at different holding potentials (as an example, a roughly hundredfold difference was found in the case of fluoxetine^2^), and a shift of the steady-state availability curve caused by drug binding is a common phenomenon. These two phenomena are not only related, but they both are manifestations of state-dependent affinity. Determining concentration-response curves at different holding potentials, and determining the shift of steady-state availability curves at different drug concentrations are essentially equivalent experiments, as it has been discussed before – see Fig. 1 of Lenkey *et al*.^1^. It is a general practice, therefore, that instead of a single IC_50_ value, the resting-state-, and inactivated-state-affinities (*K*_*R*_ and *K*_*I*_) are given for individual compounds. Once *K*_*R*_ and *K*_*I*_ are known, the potency at any membrane potential can be estimated. Excitable cells, however, do not keep a constant membrane potential, but fire action potentials regularly. Whenever an action potential is fired, sodium channels undergo a series of conformational transitions, and sodium channel inhibitors dynamically associate and dissociate depending on the actual conformational distribution of the channel population. The final effect of the inhibitor will depend on how the firing rate (the temporal pattern of the membrane potential) and binding/unbinding kinetics relate to each other. This is the basis of the well-known difference between subclasses of class I antiarrhythmics, but binding/unbinding kinetics is equally important in the therapy of hyperexcitability-related skeletal muscle disorders^3,4^, as well as diseases of the peripheral and central nervous system, such as certain pain syndromes and epilepsies. When assessing the onset/offset kinetics of a sodium channel inhibitor, one must consider the special position of the local anesthetic binding site: it is located within the central cavity of the channel, accessible only through the lipid membrane. The onset/offset process, therefore, cannot be simplified into a single-step binding/unbinding reaction^5^. The onset is often not diffusion-limited, but hindered by other possible rate-limiting steps: deprotonation of charged nitrogens (evidenced by the pH-dependence of onset rates^6^), partitioning into the membrane (evidenced by the correlation between lipophilicity and potency^1,7^), access to the central cavity through the fenestrations and the activation gate (these open up only at depolarized conformations^8^), and formation of the high-affinity binding site (the whole binding pocket is thought to be rearranged at depolarized conformations). Rate limiting steps during offset may include delayed conformational rearrangement of the protein, unbinding, egress from the central cavity, and partitioning of the drug molecule into the extracellular aqueous phase. The last process may be further delayed if the compound has accumulated within intracellular lipid compartments, the depletion of which might require a longer time.

**Fig 1.**
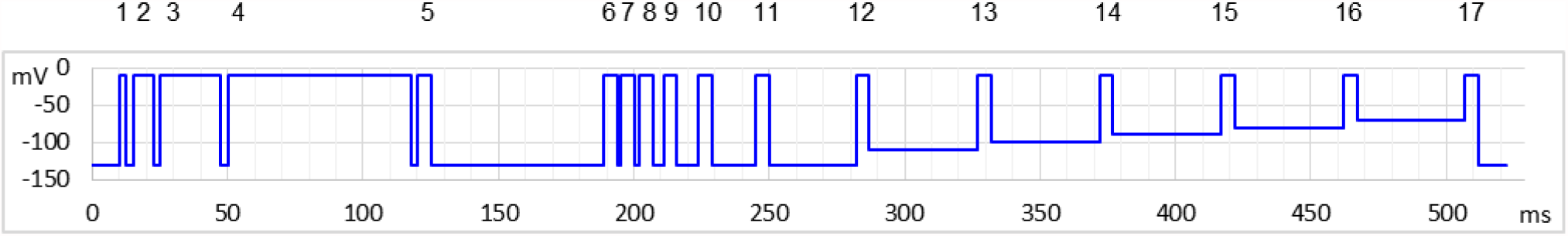
Schematic picture of the voltage protocol. Pulses are numbered for reference.

The development of the automated patch-clamp technique has made it possible to directly test the effect of multiple compounds on ion channels. However, in the case of sodium channel inhibitors, determination of an IC_50_ value, or even K_R_ and K_I_ values will not predict the therapeutic potential of specific compounds. One should achieve a comprehensive characterization of the mechanism of action for each compound. This, however, usually requires several months of experiments and analysis in a conventional manual patch clamp lab. Our aim was to design a method that could give us a detailed picture of the processes involved in the mechanism of action for individual compounds, without increasing the required time or the cost of measurements. We attempted to reconcile high throughput screening with detailed analysis of the mechanism of action, by maximizing useful information obtained during a rapid test of the compounds.

## Methods

### Cell culture and expression of recombinant sodium channels

The recombinant rNaV1.4 channel-expressing cell line was generated as described before^9^ by transfection of rNaV1.4 BAC DNA constructs into HEK 293 cells (ATCC CRL-1573) by Fugene HD (Promega, Fitchburg, WI) transfection reagent according to the manufacturer’s recommendations. Cell clones with stable vector DNA integration were selected by the addition of Geneticin (Life Technologies, Carlsbad, CA) antibiotic to the culture media (400 mg/ml) for 14 days. HEK293 cells were maintained in Dulbecco’s Modified Eagle Medium, high glucose supplemented with 10% v/v fetal bovine serum, 100 U/ml of penicillin/streptomycin, and 0.4 mg/mL Geneticin (Life Technologies, Carlsbad, CA). T175 – T25 For experiments cells were plated onto T25 (for Port-a-Patch experiments) or T175 (for IonFlux experiments) flasks, and cultured for 24–36 h. Before experiments cells were dissociated from the dish with Accutase (Corning), shaken in serum-free medium for 60 minutes at room temperature, then centrifuged, and resuspended into the extracellular solution to a concentration of 5×10^6^ cells/mL.

### Automated patch clamp electrophysiology

Ensemble voltage-clamp recordings were performed on an IonFlux Mercury instrument (Fluxion Biosciences). Cell suspension, intracellular solution, and drug-containing extracellular solution were pipetted into the 384-well IonFlux microfluidic ensemble plates. Ensemble plates are divided into four “zones”, typically each zone was used for a separate experiment (one particular set of compounds on one particular cell line). Each zone consists of 8 separate sections, which are distinct functional units, containing one well for the cell suspension, one well for the waste, two cell “traps” (intracellular solution-filled wells under negative pressure to establish high resistance seals and then whole-cell configuration), and eight compound wells. The composition of solutions (in mM) was: Intracellular solution: 50 CsCl, 10 NaCl, 60 CsF, 20 EGTA, 10 HEPES; pH 7.2 (adjusted with 1 M CsOH). Extracellular solution: 140 NaCl, 4 KCl, 1 MgCl_2_, 2 CaCl_2_, 5 D-Glucose and 10 HEPES; pH 7.4 (adjusted with 1 M NaOH). The osmolality of intra- and extracellular solutions was set to ∼320 and ∼330 mOsm, respectively. Data were sampled at 20 kHz, and filtered at 10 kHz. Experiments were carried out at room temperature.

### Single-cell electrophysiology

Port-a-Patch (Nanion, Munich, Germany) experiments were used to validate the automated patch-clamp protocol and experimental data. Whole-cell currents were recorded using an EPC10 plus amplifier and the PatchMaster software (HEKA Electronic, Lambrecht, Germany). During cell catching, sealing and whole-cell formation, the PatchControl software (Nanion) commanded the amplifier and the pressure control unit. The resistance of borosilicate chips was between 2.0 and 3.5 MΩ. The composition of solutions was identical to the ones used in IonFlux Mercury experiments.

### Rationale for the automated patch clamp voltage- and drug perfusion-protocol

In excitable cells sodium channels continuously change their conformations depending on the membrane potential. On the one hand, binding and unbinding of drugs are conformation-dependent, on the other hand, drug binding alters conformational transitions (gating) of channels. These interactions produce a special dynamics of continuously changing drug potency: it does not only depend on the actual value of membrane potential, but also on its recent history. To assess both membrane potential dependence and time dependence, we used the protocol illustrated in Fig. 1. We choose to study three aspects of membrane potential-dependent dynamics of drug potency: First, the effect of inhibitors often needs some time to develop. In the first section of the protocol (pulse #1 to #5), therefore, we intended to assess how fast the effect of the drug develops upon depolarization. We used progressively lengthened depolarizations and monitored the inhibition. Second, inhibitors most often dissociate from hyperpolarized (resting) channel conformation, therefore, drug potency gradually decreases upon prolonged hyperpolarization. In the second section of the protocol (pulses #6 to #12) we assessed the dynamics of this recovery using progressively lengthened hyperpolarizations. Third, we assessed quasi-equilibrium conditions: we investigated in this section (pulses #13 to #17) how the extent of inhibition depended on the membrane potential. The three sections of the protocol correspond with the protocols “state-dependent onset” (**SDO**), “recovery from inactivation” (**RFI**), and “steady-state inactivation” (**SSI**) we used in previous studies^9,10^, although with some significant differences. Our priority with this current protocol was high time resolution.

For this reason, the duration of the whole 17-pulse protocol was only 522 ms, and it was repeated every second throughout the experiment. A standard experiment included 7 different drug applications, 40 s long each, with 60 or 80 s wash periods between them, then the whole sequence was repeated. This means that the experiment lasted for 28-30 min, during which ∼1700-1800 sweeps were recorded.

The microfluidic plate used in experiments contains 8 compound wells, thus it would allow perfusion of 8 different compounds. However, we found that solution exchange was faster and more reliable if we used compound well #1 to perfuse control extracellular solution throughout the experiment.

In the **SDO** section of the protocol, we tested the effect of only four depolarization durations (besides the control): 2.5, 7.5, 22.5, and 67.5 ms. We used a cumulative arrangement, not allowing full recovery between depolarizations (only 2.5 ms at hyperpolarized potential between depolarizations). We used Port-a-Patch experiments (i.e., in gigaseal, single-cell recordings) to verify the effects observed in IonFlux experiments (i.e., in multi-cell recordings with varying seal resistance); and also to compare the effect of this cumulative arrangement of the protocol with the conventional multi-sweep protocol, where all sweeps are started with the whole channel population in resting state. Protocols similar to this one are often used to study slow inactivation. It is important to note that in our experiments the **SDO** protocol was not intended for the study of slow inactivation, but the study of drug effect onset, upon depolarization-induced conformational change. Depolarized conformations (open and inactivated) provide increased affinity, and/or increased accessibility to the binding site, thereby allowing the development of a new binding/unbinding equilibrium. The protocol investigates how fast this new equilibrium is reached. Slow inactivation may only play a minor role in the development of the effect, since even the longest duration (64 ms) is insufficient to induce substantial slow inactivation. The interpulse interval (2.5 ms) was chosen so that it would not allow full recovery even from fast inactivation. This way we maximized sensitivity to drug effects: if any drug stayed bound for at least 2.5 ms then it produced either channel block or delayed recovery by modulation; in both cases, the effect was sure to be detected. Thus far we have encountered only one single compound that could fully dissociate within 2.5 ms, and thus its effect was undetected in the **SDO** protocol (see the accompanying paper^11^).

In the case of the **RFI** protocol, we used 8 hyperpolarization durations: 1, 2, 4, 8, 16, 32, 64, and 498 ms. The 64 ms hyperpolarization also served to separate the **SDO** and **RFI** sections of the protocol (to allow time for recovery). In the case of slowly acting drugs, where state-dependent binding equilibrium was not reached within 64 ms, we occasionally observed non-monotonous recovery (see the legend of Fig. S1 for discussion). The longest (498 ms) hyperpolarization was not recorded (except its first 10 ms after the last pulse and its last 10 ms before the first pulse of the next sweep).

Instead of a conventional **SSI** protocol, where full recovery to resting state is allowed between sweeps of the protocol, we used an accelerated procedure to assess membrane potential dependence, which did not include hyperpolarizations. This means, that resting/inactivated equilibrium was approached from a fully inactivated channel population, not from a fully resting population. If there was a true steady-state, this would make no difference. We used 40 ms pre-pulse duration, which allowed ∼90% recovery from fast inactivation at −130 mV membrane potential, and somewhat less at less negative potentials. Consequently, this “no-hyperpolarization” protocol gave similar results to the conventional protocol under control conditions (see Fig. S2), but was more sensitive to drug effects: it detected a larger V_1/2_ shift because the effect of shifted equilibrium was accompanied by the effect of delayed recovery from inactivation. It also detected a larger delay of recovery from inactivation. The extent of difference depended on drug onset/offset dynamics. Beyond rapidity of testing, there was an additional practical advantage of the “no-hyperpolarization” protocol: It eliminated the problem of sub-threshold activation during pre-pulses due to poor space clamp. This is important in automated patch clamp instruments which record from a group of cells, and therefore seal resistance and series resistance values for individual cells are poorly controlled.

Major properties of gating kinetics and equilibrium in control, as well as in the presence of lidocaine and riluzole have been repeated in Port-a-Patch experiments to assess the quality of measurements in the IonFlux Mercury instrument and to compare conventional and cumulative protocols. The results are shown in the Supplement (Fig. S2).

### Data analysis

Using the data acquisition software, all data traces were exported as CSV files. A custom software was developed in Octave to automatically process raw data. All traces were read in from the DataAcquisition* .ISD files containing the raw data. Separate csv files contained the description of the voltage protocol. First, 2 ms sections were selected after each of the 34 voltage steps. Capacitive artifacts were removed by calculating the sum of the section recorded at the beginning and after the end of the pulse, and then subtracting the offset so that all currents started at zero current level. Although other voltage-gated channels were present in the cells, their contribution to the fast transient inward current was small (the peak amplitude of the TTX-resistant fraction of the fast transient inward current was 5.17 ± 3.44% of the full amplitude), thus allowing reasonably accurate assessment of the extent of sodium channel inhibition. For all ∼1700 sweeps, and for all 17 pulses, the minima (peak amplitudes) were extracted and saved in 64 csv files for the 64 cell ensembles. These data (all 17 peak amplitudes for each second of a ∼1700s experiment plotted against time) are shown for one particular cell ensemble in Fig. 2. From each of the four zones (separate experiments), we chose n = 6 ensembles for analysis, based on the stability of amplitude and seal resistance throughout the experiment.

**Fig 2.**
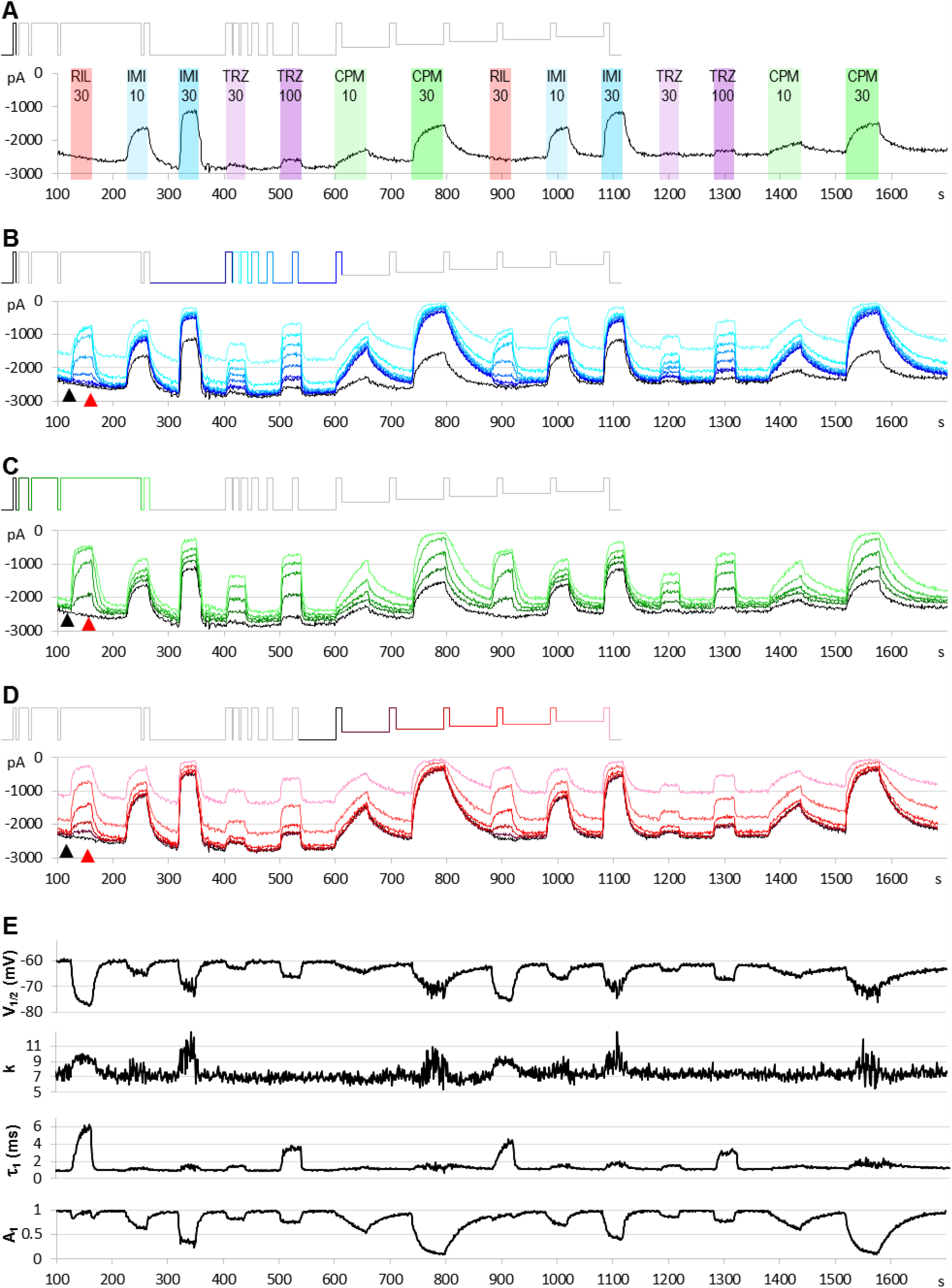
An example for the plot of current amplitudes throughout the experiment. The experiment included two repetitions of seven different compound applications: 30 µM riluzole, 10 and 30 µM imipramine, 30 and 100 µM trazodone, 10 and 30 µM chlorpromazine. Currents evoked by all 17 depolarizations are shown, grouped into three functional assays, as described in the text. (A) Peak amplitude plot for pulse #1-evoked current throughout the experiment. (B) Currents evoked by pulses #1 and #6-#12; these allow second-to-second reconstruction of recovery from inactivation (RFI; see Methods) throughout the experiment. (C) Currents evoked by pulses #1-#5; these allow reconstruction of the **SDO** plots (see Method). (D) Currents evoked by pulses #1 and #13-#17; these allow reconstruction of the **SSI** plots (see Method). Insets show the schematic picture of the voltage protocol, where colors match the color of the corresponding amplitude plot. (E) Results of the automated analysis. Automated fitting of **SSI** and **RFI** data was done as described in Methods. The plot shows the changes in half inactivated voltage (*V*_*1/2*_) and slope (*k*) values from **SSI** data, and the value (*τ*_*1*_) and contribution (*A*_*1*_) of the fast time constant from RFI data.

**SSI** and **RFI** plots for each sweep were fitted by the Octave script. Fitting 64 × 1700 plots could not be individually visually supervised, but parameters of the automated fitting were evaluated by comparing them to visually controlled fits of **SSI** and **RFI** plots, as shown in Results. At the end of all drug perfusion periods, as well as at the end of control periods before and after them, we fitted data using the Solver add-in of Microsoft Excel. The precision of the automated fit and the adequacy of the equation used were evaluated; if we found the fitting inadequate, either the equation or the constraints were modified. To evaluate the precision of the fit we calculated the root mean square error (*RMSE*) values for each fit, as well as relative error (*E*_*rel*_) values for each point, were recorded and saved in a separate csv file. The following formulas were used to calculate the extent of error:

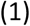

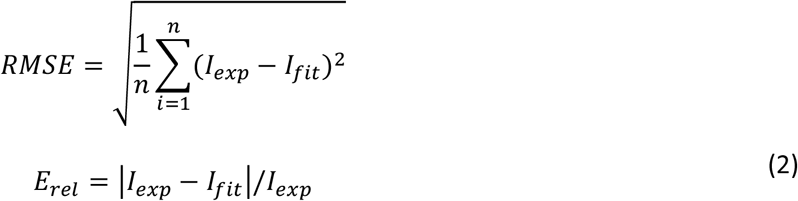

where *n* is the number of points fitted, *I*_*exp*_ is the experimentally measured amplitude, and *I*_*fit*_ is the fitted amplitude. We expressed *RMSE* values as a percentage of the maximal amplitude (for each sweep, the amplitude of the current evoked by pulse #1 of that particular sweep). The advantage of automated fitting of all **SSI** and **RFI** plots throughout the experiment was, that second-to-second changes in *V*_*1/2*_, *k, A*_*1*_, and *τ* values reveal the dynamics of development/removal of modulatory drug effect more accurately; furthermore, even minimal effects were detectable, because the tests were repeated several times before, during, and after drug applications.

Peak amplitudes of currents evoked by pulses #12 to #17 were used to construct **SSI** curves, which were fit using the Boltzmann function:

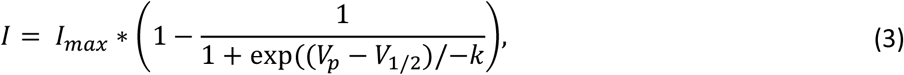

where *V*_*p*_ is the pre-pulse potential, *V*_*1/2*_ is the voltage where the curve reached its midpoint and *k* is the slope factor. Currents evoked by pulses #1, and #6 to #12 were used to construct **RFI** plots, which were fitted with a bi-exponential function:

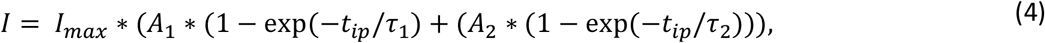

where *τ*_*1*_ and *τ*_*2*_ are the fast and slow time constants, *A*_*1*_ and *A*_*2*_ are their respective contribution to the amplitude, and *t*_*ip*_ is the duration of the interpulse interval. We routinely used constrains *τ*_*1*_ < *τ*_*2*_, and *A*_*1*_ + *A*_*2*_ = 1, and for automated fitting we also constrained the slow time constant. We found that in the presence of riluzole equation (4) could not adequately fit RFI plots, therefore we used an extended equation:

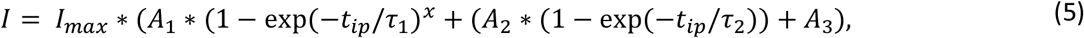

where *A*_*3*_ represents the unmodulated fraction of channels at low riluzole concentrations which recover as control channels, and the exponent “*x*” was needed because recovery in the presence of riluzole has been repeatedly found to be steeper than exponential. (This has been addressed either by including a time delay parameter in the equation^12,13^, or by using an equation where the fast exponential component was on the *x*^*th*^ power^9,14^. We prefer the latter, because introducing a delay parameter results in negative numbers at short time intervals.)

We did not perform an automated fit of **SDO** plots, because fitting often required different functions for different drugs. Analysis of **SDO** data is described in the accompanying paper^11^.

The microfluidics of the IonFlux instrument could provide complete solution exchange within the 498 ms hyperpolarization (478 ms of which was unrecorded), therefore solution exchange rate did not compromise kinetic analysis of data. Complete solution exchange between sweeps was verified using high Ca^2+^ ion concentration (35 mM) containing solution (which blocks sodium channels).

In a regular experiment, only the first set of compound applications were evaluated, repetition of the experiment served as an internal control: it helped to detect incomplete recovery (see *e*.*g*. after 30 µM chlorpromazine in Fig. 2), and to verify onset and offset rates (see *e*.*g*. the offset after 30 µM imipramine, where some disturbance obscured the offset process). It also helped to assess the extent of the spontaneous leftward shift of the steady-state availability curve by observing the ratio of the 17^th^/12^th^ pulse evoked current amplitudes (pink and darkest red traces in Fig. 2D, 4A, and 5A).

## Results

### Initial examination of data

In order to better explain how to interpret our data, we will first show an example for a single experiment, and make a few important general observations. After explaining the interpretation of results, we will describe how quantitative analysis from multiple experiments with multiple concentrations is performed, on the example of lidocaine and riluzole. Figure 2 illustrates the results of a single experiment. In this example, we perfused the following compounds: riluzole (30 µM), imipramine (10 and 30 µM), trazodone (30 and 100 µM), and chlorpromazine (10 and 30 µM). The full voltage protocol (Fig. 1) is described in the Methods section, in the interest of clarity here we will discuss it as if it was built up step-by-step.

Let us first consider what would happen if we gave only single depolarizing pulses at every second (Fig. 2A). The peak amplitude plot shows that the amplitude was fairly stable throughout the ∼30-minute experiment. Riluzole at 30 µM caused no inhibition whatsoever, trazodone inhibited peak amplitudes only minimally (∼5% inhibition at 100 µM), the other two compounds caused concentration-dependent inhibition. Imipramine seemed to be the most potent compound, causing ∼50-55 % inhibition at 30 µM.

Let us now consider the section of the experiment in which we tested the rate of recovery from inactivation (**RFI**). Current amplitudes evoked by the highlighted part of the voltage protocol are plotted throughout the experiment in Fig. 2B. Colors in the voltage protocol match colors in the current amplitude plot. We applied consecutive depolarizing pulses with increasing interpulse intervals between them within a single sweep (*i*.*e*., within a single uninterrupted period of data acquisition). Interpulse intervals were 1, 2, 4, 8, 16, 32, 64, and 498 ms (not in this sequence, see colors in the protocol, as described in more details in Methods). Peak amplitudes evoked after 1 ms hyperpolarization are shown as a light blue line, currents evoked after progressively longer interpulse intervals are shown by increasingly darker shades of blue. The black line indicates the current evoked after 498 ms hyperpolarization, it is identical to the one shown in Fig. 2A. We can observe that, in contrast to what we saw in Fig. 2A, riluzole (30 µM) and trazodone (100 µM) did produce a massive inhibition, only the inhibition by these compounds was transient, re-appearing and disappearing within each 1 s cycle. We can observe in the case of riluzole that inhibition already started to ease off at the 4 ms interpulse interval, and it almost completely disappeared by the end of the 16 ms interpulse. Inhibition by trazodone disappeared incrementally, some residual inhibition was present even at 498 ms. In contrast, inhibition by imipramine or chlorpromazine recovered minimally within 64 ms, substantial recovery only occurred during the longest (498 ms) interpulse interval. Note that two fundamentally different processes can be observed in Fig. 2B: One can discern a dynamics of onset and recovery within individual sweeps, on a millisecond time scale (see Fig. 3, below), in the continuous presence of the drug; we will call this “micro-dynamics”. The dynamics of onset and offset upon drug application and removal, on the other hand, occurred on the time scale of seconds, we will call that “macro-dynamics”. Macro-dynamics provides valuable information regarding the physicochemical properties of individual compounds, which determine their *in vivo* pharmacokinetics, and the extent of their accumulation within the plasma membrane and intracellular compartments. Note, however, that it is micro-dynamics that determines firing frequency-dependent inhibition of excitable tissues. Macro-dynamics, as observed upon rapid wash-in and wash-out of drug-containing solution *in vitro*, does not occur during *in vivo* drug delivery.

**Fig 3.**
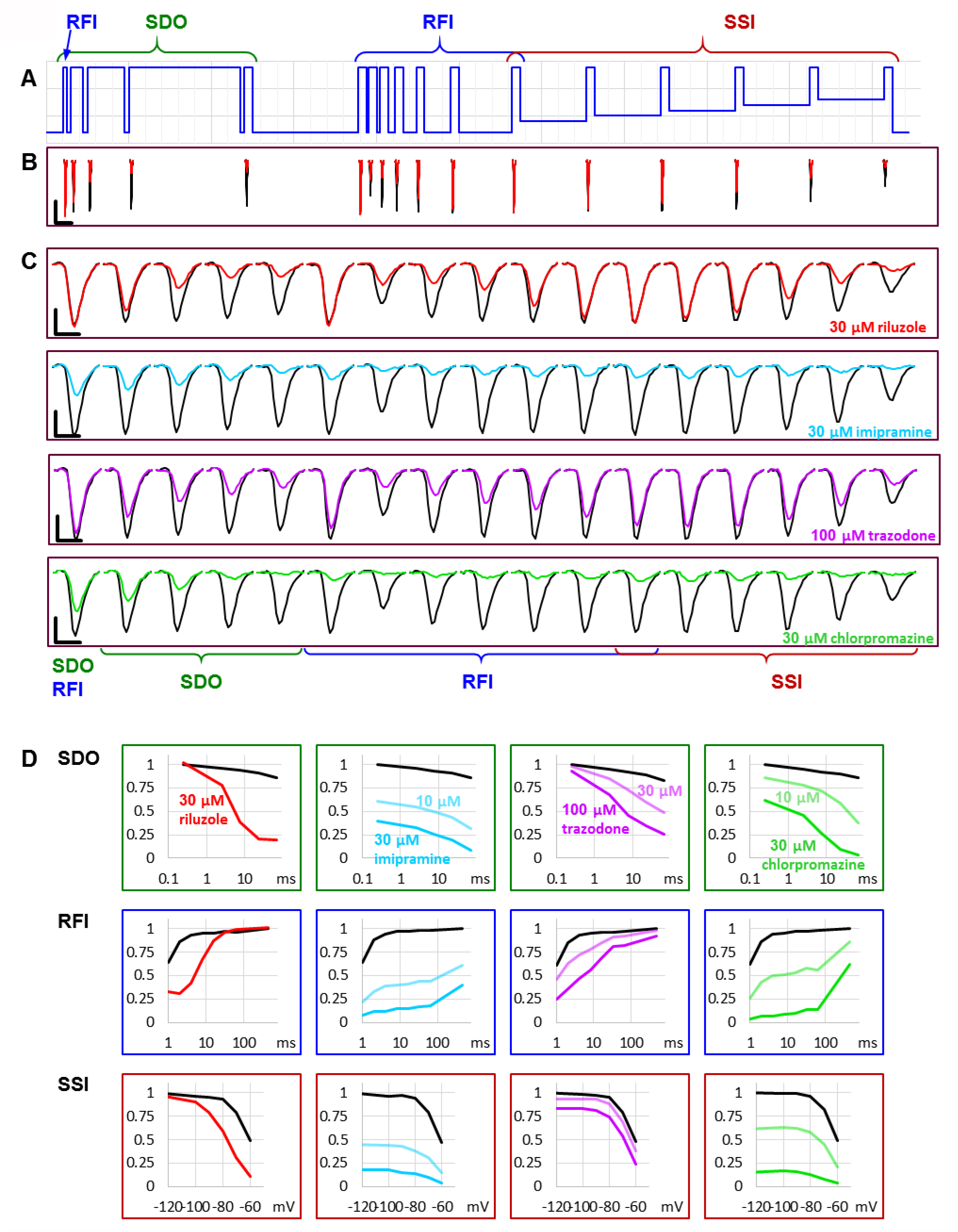
Reconstruction of SDO, RFI, and SSI plots from the current amplitude plots. (A) The voltage protocol (for reference). (B) Evoked currents in control, and in the presence of 30 µM riluzole, shown on the same time scale as the scheme of the protocol. Note the difference in the potency of riluzole between subsequent depolarizations. Scale bars: 1 nA, 10 ms (C) Evoked currents on an expanded time scale for visibility, before, and during the perfusion of riluzole, imipramine, trazodone, and chlorpromazine. Scale bars: 1 nA, 1 ms (D) Reconstruction of **SDO, RFI**, and **SSI** plots before, and during the perfusion of the indicated compounds.

Next, let us observe the section of the protocol which investigates “state-dependent onset” (**SDO)**, *i*.*e*., the micro-dynamics of inhibition onset at depolarized membrane potential (Fig. 2C). It shows currents evoked by depolarizations #1 to #5, as shown by the colors of the highlighted section of the protocol. We can observe that the micro-dynamics of onset can be rather different from that of recovery, for example in the case of chlorpromazine, we see a gradual onset during the 2.5 ms to 67.5 ms depolarizations, while we can observe that most of the recovery occurred between 64 and 498 ms.

Finally, depolarizations #12 to #17 assess steady-state availability at membrane potentials −130, −100, −90, −80, −70 and - 60 mV (Fig. 2D)., showing depolarizations; different shades of red).

Parameters of the automatized fitting of **SSI** and **RFI** plots are shown in Fig. 2E. We can observe again that different compounds behaved differently. Riluzole, which seemed to have no effect at all in Fig. 2A, was the most potent of all drugs in terms of shifting the V_1/2_ value. Trazodone was the only compound that did not affect the slope of the availability curve (*k*). Delayed recovery from inactivation in the case of riluzole and trazodone was predominantly due to an increase in the fast time constant, while in the case of imipramine and chlorpromazine, the fast time constant was unchanged, but its contribution was decreased. The accuracy of fits throughout the experiment can be monitored by automated calculation of *RMSE* and *E*_*rel*_ values (Fig. S1)

### Let us call attention to a few important points

Note that some of the inhibitors cause widely different extents of inhibition, depending on which of the 17 traces we consider. Figure 3 illustrates micro-dynamics that took place within a single 522 ms long sweep. The voltage protocol is shown again for reference in Fig. 3A, evoked currents are illustrated on the same time scale in Fig. 3B, right before the first application of 30 µM riluzole (black traces), and at the end of riluzole perfusion (red traces). Black and red triangles in Fig 2 indicate the exact time of the sweep from which original currents were taken. Figure 3C shows currents on an expanded time scale, we illustrate micro-dynamics during the perfusion of 30 µM riluzole (red traces), 30 µM imipramine (blue traces), 100 µM trazodone (purple traces), and 30 µM chlorpromazine (green traces). Conventional plots of **RFI, SDO**, and **SSI** (see *e*.*g*. ^9,10^) with the four compounds are shown in Fig. 3D.

Note also in Fig. 2, that the observed macro-dynamics (onset time constants upon drug application and offset time constants upon washout) can also be different depending on which of the 17 pulse-evoked currents we monitor. Let us consider for example the **SDO** section of the protocol (Fig. 2C) during the onset and offset of inhibition by 30 µM chlorpromazine. Channels activated by pulses #1 (amplitudes plotted in black) to #5 (amplitudes shown in light green) encounter the same exact concentration of the same compound during drug perfusion and subsequent washout. We know from calibration experiments that solution exchange in the extracellular aqueous phase is complete between two sweeps, however, the buildup and depletion of drug concentration within the membrane phase can be much slower, and it is the intramembrane concentration that the channel can perceive (contribution of the hydrophilic pathway is probably negligible for these strongly lipophilic compounds). The difference in potency and dynamics between traces (e.g. the light green trace and the black trace in Fig. 2C) reflects different sensitivities of the channel, depending on its recent gating history.

If we compare the pattern produced by 100 µM trazodone and 30 µM chlorpromazine, we can observe an obvious difference not only between their macro-dynamics (both onset and offset were clearly slower for chlorpromazine) but also between their micro-dynamics. In both cases we see a gradually deepening inhibition in the **SDO** section, indicating that the onset of inhibition occurred within the investigated time window (2.5 to 67.5 ms). Recovery, however, was different: the effect of trazodone recovered rapidly, mostly within 64 ms (see pulse #6 in Fig. 3C); while inhibition by chlorpromazine was not much relieved throughout the whole 17-pulse sweep, and substantial recovery only occurred during the 498 ms inter-sweep intervals.

Riluzole and trazodone showed intensive micro-dynamics: during the inter-sweep intervals, much of the inhibition was relaxed, while it was repeatedly re-established upon depolarizations. Imipramine, in contrast, showed minimal micro-dynamics, once the inhibition was established (by the end of the **SDO** section), the hyperpolarizations within the sweep (up to 64 ms) were not long enough to allow significant recovery. Even the 498 ms inter-sweep hyperpolarization was enough only for partial recovery. For this reason, **SSI** data could not be correctly measured, much longer periods would be required for establishing equilibrium. This protocol was optimized for the study of compounds with fast micro-dynamics, therefore it was inaccurate for slower micro-dynamics compounds. Chlorpromazine was similar to imipramine, with somewhat faster micro-dynamics, but slower macro-dynamics. A larger fraction of channels recovered during inter-sweep intervals, but during each sweep, after the inhibition was re-established (by the end of pulse #5), all pulses were inhibited similarly, because of the insufficient time for equilibration.

From the initial examination of this single experiment, it is apparent that different drugs have their own characteristic “signature” pattern of inhibition. This is evident from the similarity of repeated drug application effects, as well as from the effect of different concentrations of the same drug. When different concentrations of the same compound were applied, we could observe different extents of inhibition, different onset rates, but the offset rates were similar, and there was a uniform overall pattern (*i*.*e*. which of the 17 pulse-evoked currents were affected to what extent). It is clear that the voltage protocol we used could only appropriately characterize drugs with fast micro-dynamics; this protocol was intended to characterize compounds that could selectively inhibit pathological high-frequency firing. Similar protocols, with longer hyperpolarization and depolarization durations (and, therefore, necessarily with less temporal resolution) can be used for compounds with slower micro-dynamics.

In summary, initial examination indicated that these four compounds acted in four different ways. In the next section, we will show an example for an initial analysis of the effect of two well-known drugs, lidocaine, and riluzole in different concentrations. The accompanying paper^11^ will discuss how to derive compound-specific biophysical properties from this initial analysis.

#### Quantitative analysis

We illustrate quantitative analysis in the case of two well-known sodium channel inhibitors, lidocaine (30, 100, 300, and 1000 µM; Fig. 4), and riluzole (10, 30, 100, and 300 µM; Fig. 5). Fig. 4A and 5A show an example of the effect of both drugs on all 17 pulse-evoked current amplitudes. From the 17 peak amplitudes of each sweep, **SDO, RFI**, and **SSI** plots were reconstructed, but only **RFI** and **SSI** plots were fitted. Two different methods were used for fitting: automated fitting was performed for each sweep of the ∼1700 sweep experiment, and to validate the (uncontrolled) automated fitting, we fitted **RFI** and **SSI** plots with visual control only for pairs of control and drug-treated cell ensembles, as marked by the arrowheads in Fig. 4A and 5A. For all visually controlled fits three consecutive sweeps were averaged; the last three before each drug perfusion period, and the last three at the end of each drug perfusion. The **SDO, RFI**, and **SSI** plots, constructed from these three-point averages for six cell ensembles as well as the average of the six measurements (dashed lines)are shown in Fig. 4C and 5C. Light to dark color of plots (teal for **SDO**, indigo for **RFI**, and purple for **SSI** plots) indicate increasing concentration, these colors match the colors of corresponding arrowheads in Fig. 4A and 5A, as well as the colors of columns in Fig. 4D and 5D, where parameters from the visually controlled fitting are summarized. Six parameters are shown, *τ*_*1*_, *τ*_*2*_, *A*_*1*_, and *A*_*2*_ values for **RFI** fits, *V*_*1/2*_, and *k* for **SSI** plots. In the automated fitting procedure, the slow time constant (typically between 100 to 400 ms) was fixed, because there were few data points in this time range. We calculated the mean slow time constant from the visually controlled fitting and then used this fixed value throughout the automated fitting procedure. In addition, we used the constrain of *A*_*1*_ + *A*_*2*_ = 1. For this reason, only four parameters are shown, *τ*_*1*_, and *A*_*1*_ values for **RFI** fits (blue dots), *V*_*1/2*_, and *k* for **SSI** plots (pink dots). (Fig. 4B and 5B). For the sake of comparison, parameters obtained from the visually controlled fitting are also shown in these figures, as large circles. We can observe that the automated fitting procedure quite reliably reproduced data from the visually controlled fitting, except in the case of 10 µM and 30 µM of riluzole (see below for an explanation). The overall quality of the fit could be monitored by observing the % *RMSE* values; from the E_*rel*_ values, we could see which particular point contributes most to the error. In the case of both riluzole and lidocaine E_*rel*_ values were quite low, except for pulses #7, #8, and #17; where peak amplitudes were the smallest. Small amplitudes necessarily result in a higher relative error, both because of the decreased signal-to-noise ratio and because the fitting procedure minimizes absolute, not relative, squared errors.

**Fig 4.**
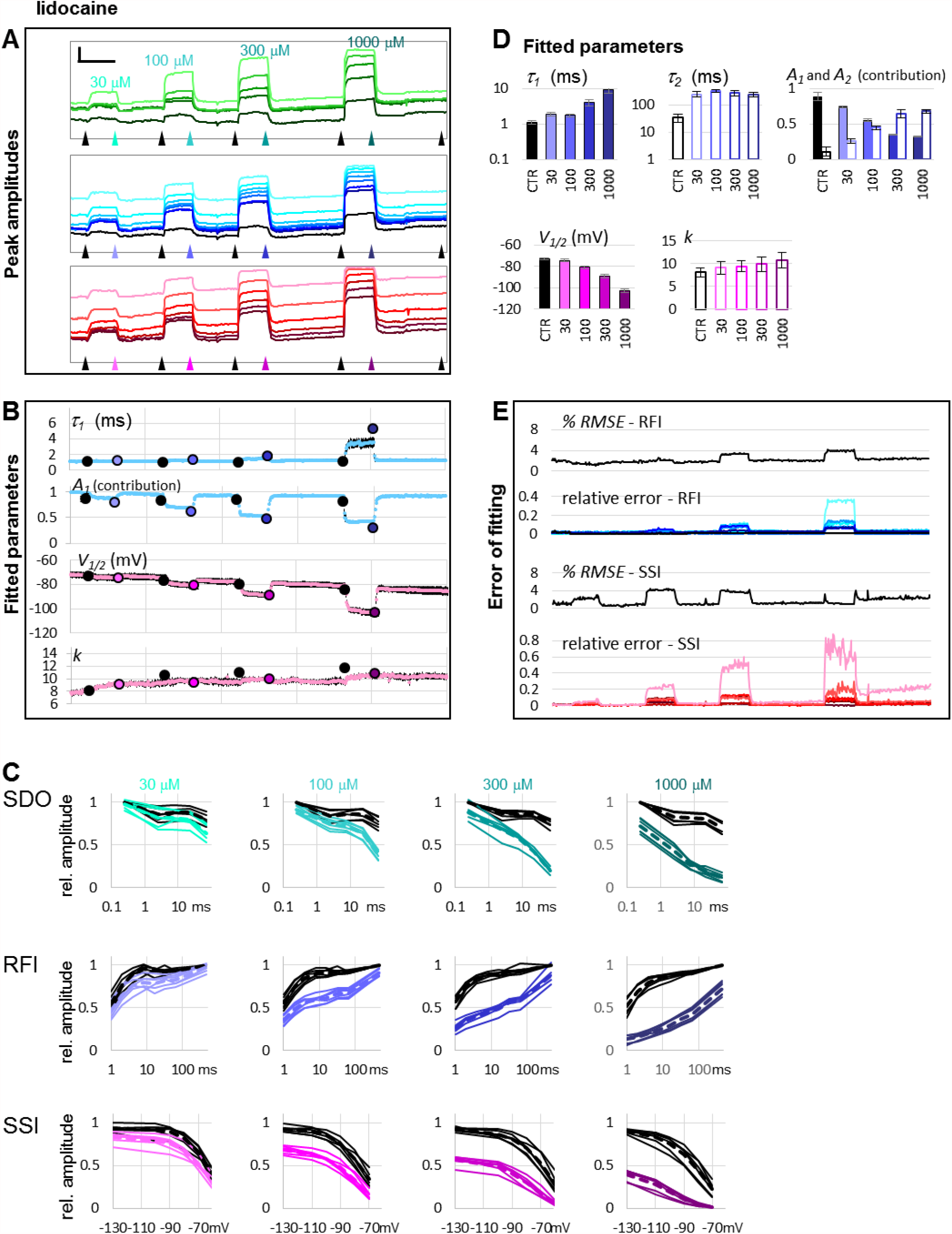
The effect of four different concentrations of lidocaine on peak amplitudes. (A) An example of peak amplitude plots for all 17 pulse-evoked currents, grouped into the **SDO, RFI**, and **SSI** groups as shown in Fig. 2. Scale bars: 2 nA, 50 s. Arrowheads indicate the time points immediately before, and at the end of drug perfusion periods. At each point, indicated by the arrowheads, the average of three consecutive data points was taken to construct the plots shown in panel C. (B) Parameters of Boltzmann, and bi-exponential automated fits throughout the four drug application periods. Colored circles indicate the values obtained by visually controlled fitting of n = 6 curves; the same data that is shown in panel C. (C) Reconstructed **SDO, RFI**, and **SSI** plots for n = 6 cell ensembles. Colors match the colors of arrowheads in panel A. Dotted lines show the average of the six cell ensembles. (D) Parameters obtained by visually controlled fitting of RFI and SSI plots. (E) The error of fitting throughout the four drug application periods. *RMSE* values are expressed as the percentage of the peak amplitude evoked by pulse #1 of the same sweep. Relative errors are shown for pulses #6 to #12 and #1 (shades of blue and black, as shown in panel A), as well as for pulses #12 to #17 (shades of red, as shown in panel A).

**Fig 5.**
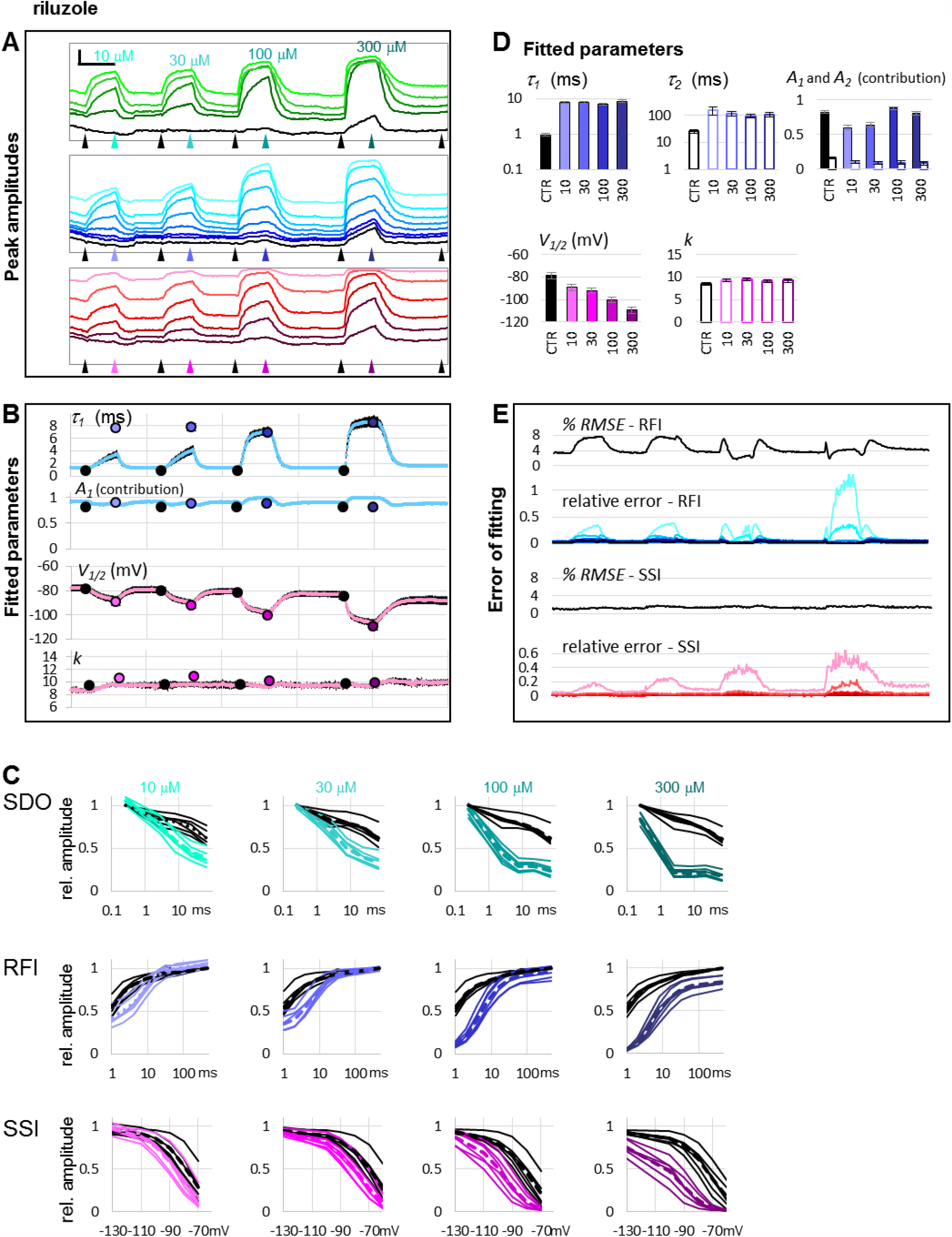
The effect of four different concentrations of riluzole on peak amplitudes. (A) An example of peak amplitude plots for all 17 pulse-evoked currents, grouped into the **SDO, RFI**, and **SSI** groups as shown in Fig. 2. Scale bars: 4 nA, 50 s. Arrowheads indicate the time points immediately before, and at the end of drug perfusion periods. At each point, indicated by the arrowheads, the average of three consecutive data points was taken to construct the plots shown in panel C. (B) Parameters of Boltzmann, and bi-exponential automated fits throughout the four drug application periods. Colored circles indicate the values obtained by visually controlled fitting of n = 6 curves; the same data that is shown in panel C. (C) Reconstructed **SDO, RFI**, and **SSI** plots for n = 6 cell ensembles. Colors match the colors of arrowheads in panel A. Dotted lines show the average of the six cell ensembles. (D) Parameters obtained by visually controlled fitting of RFI and SSI plots. For RFI plots Eq #5 was used, with the constraint of *x* = 1. (E) The error of fitting throughout the four drug application periods. *RMSE* values are expressed as the percentage of the peak amplitude evoked by pulse #1 of the same sweep. Relative errors are shown for pulses #6 to #12 and #1 (shades of blue and black, as shown in panel A), as well as for pulses #12 to #17 (shades of red, as shown in panel A).

In the **RFI** plots, lidocaine caused a slowing of the fast time constant of recovery, which was moderate at lower concentrations (30 and 100 µM) but was substantial at 100 and 300 µM concentrations (∼4-fold and ∼7-fold, respectively). More importantly, the contribution of the slow time constant gradually overcame the contribution of the fast one, increasing from 8.8 ± 2.1 % at control, to 68.8 ± 1.4 % at 1000 µM, while the value of the slow time constant did not change with concentration (Fig. 4D). In contrast, riluzole caused no change in either the contribution or the value of the slow time constant but caused a radical ∼8-fold increase in the fast time constant. The extent of this increase did not change significantly with concentration (ranging from 7.4-to 8.9-fold), but at lower concentrations there seemed to be an unmodulated fraction of the channel population, as it can be seen on the contribution of *A*_*1*_ at concentrations 10 and 30 µM (Fig. 5D).

In the **SSI** plots, both compounds caused a concentration-dependent hyperpolarizing shift in the half inactivation voltage, while the slopes did not change significantly.

In summary, it was possible to perform automated fitting of ∼1700 **RFI** and **SSI** plots on 64 channels, with reasonable accuracy. *RMSE* values remained below 4%, except for **RFI** plots in the presence of riluzole, where the conventional bi-exponential equation was clearly inadequate for fitting the data. In this special case recovery data have been consistently found to be steeper than exponential, and therefore either have been fit with an equation that included a time delay parameter^12,13^, or with an equation where the fast exponential component was on the *n*^*th*^ power^9,14^.) Visually controlled fits confirmed the parameters of automated fitting and gave somewhat better *RMSE* values (between 0.44 and 1.56 % for all **SSI** fits, and between 1.12 and 2.27 % for **RFI** fits in control, and in the presence of lidocaine). In the case of **RFI** plots in the presence of riluzole, visual control allowed us to identify the source of error, and to modify the equation accordingly. Fitting the **RFI** plot with the simple bi-exponential equation gave *RMSE* values 3.54, 2.97, 1.79, and 2.76 %, for 10, 30, 100, and 300 µM riluzole, respectively. To improve these, we introduced the extended equation (Eq. 5) in two steps: in the first step, we allowed *A*_*3*_ to be different from zero. Allowing a non-zero unmodulated fraction was important in the case of 10 and 30 µM concentrations, and improved their *RMSE* values to 0.64 and 0.67, respectively, but did not change the *RMSE* values for 100 and 300 µM riluzole. The parameters obtained with this modification are shown in Fig. 4D. In the next step, we also allowed the exponent *“x”* to be different from 1. This improved *RMSE* values for all four concentrations: to 0.50, 0.64, 1.04 and 1.11 % for 10, 30, 100, and 300 µM riluzole, respectively. By allowing the exponent to vary, however, we lost the comparability of fast time constants, because the time constant and the exponent are interdependent, as we have discussed before^9^. This is the reason why in Fig. 4D we show time constants from the fit when non-zero *A*_*3*_ of Eq. 5 was allowed, but the exponent was not allowed to differ from 1.

## Discussion

Our first important observation is the obvious existence of two completely different processes, named micro- and macro-dynamics. This emphasizes the often unappreciated complexity of the processes which underlie the onset and offset of drug effect. Macro-dynamics is often studied using single pulses delivered at a certain fixed frequency, while the compound is washed in and then washed out. Macro-onset and macro-offset time constants are commonly interpreted as reflecting association and dissociation, and therefore are used to determine the affinity of binding. However, these processes never occur *in vivo*, and most likely do not reflect purely association and dissociation (only if partitioning into the membrane and access into the central cavity are relatively unobstructed, and therefore the rate-limiting step of onset is diffusion itself).

In contrast, micro-dynamics keeps going on incessantly all the time, in all excitable cells of the organism. Micro-offset is conventionally studied using the recovery from inactivation (RFI) protocol (using different interpulse intervals in separate sweeps). This recovery is also commonly interpreted as dissociation. Therefore, it is important to point out that for many inhibitor compounds micro- and macro-dynamics differ by several orders of magnitude, therefore it is not clear whether unbinding itself contributes to one, to the other, or both. For most compounds, we suppose that macro-onset may reflect deprotonation (as evidenced by the pH dependence of macro-dynamics^6^), accumulation within the membrane phase, and state-dependent access to the central cavity. Macro-offset may reflect state-dependent egress from the central cavity and depletion of the membrane phase (and intracellular compartments). Micro-dynamics, for most inhibitors, probably reflects genuine binding/unbinding, but may also represent modulated gating^14^ or the process of access/egress (diffusion between the central cavity and the membrane phase). Micro-dynamics is always dependent on conformational states, and in turn, it also affects the distribution of conformational states (by altering the rates of transitions between them).

Interestingly, although micro-dynamics obviously cannot be slower than macro-dynamics, the two are in fact not strongly correlated. We have found compounds with fast micro-dynamics but relatively slow macro-dynamics (e.g. riluzole, with more than 1000-fold difference between micro- and macro-dynamics rates), and also some with relatively slow micro-but fast macro-dynamics (like bupivacaine, with less than 10-fold difference).

For slow micro-dynamics compounds, we will need to prolong the experimental protocol with both longer depolarizations and hyperpolarizations. Of course, this can only be done at the expense of time resolution, but for drugs, with slow dynamics, this seems acceptable. Allowing close-to-full equilibration of micro-onset and micro-offset should not compromise macro-dynamics.

In summary, automated analysis of data obtained using a complex voltage- and drug application protocol for an automated patch clamp instrument allowed us to assess the complex dynamics of drug-ion channel interaction, and to identify multiple sub-processes that constitute the onset and offset of drug effects during and after drug perfusion. Dynamics of drug binding/unbinding, access/egress, protonation/deprotonation, as well as the extent of modulation and channel block are crucial determinants of therapeutic effectiveness. We expect that a comprehensive assessment of the mechanism of action can provide a better prediction of therapeutic potential than the assessment of resting and inactivated affinity only.

## List of non-standard abbreviations

SDO: “state-dependent onset” protocol
RFI: “recovery from inactivation” protocol
SSI: “steady-state inactivation” protocol
*%RMSE*: percentage root mean square error
*E*_*rel*_: relative error
EIP: effective inhibitor potency

## Supporting information

**Fig S1.**
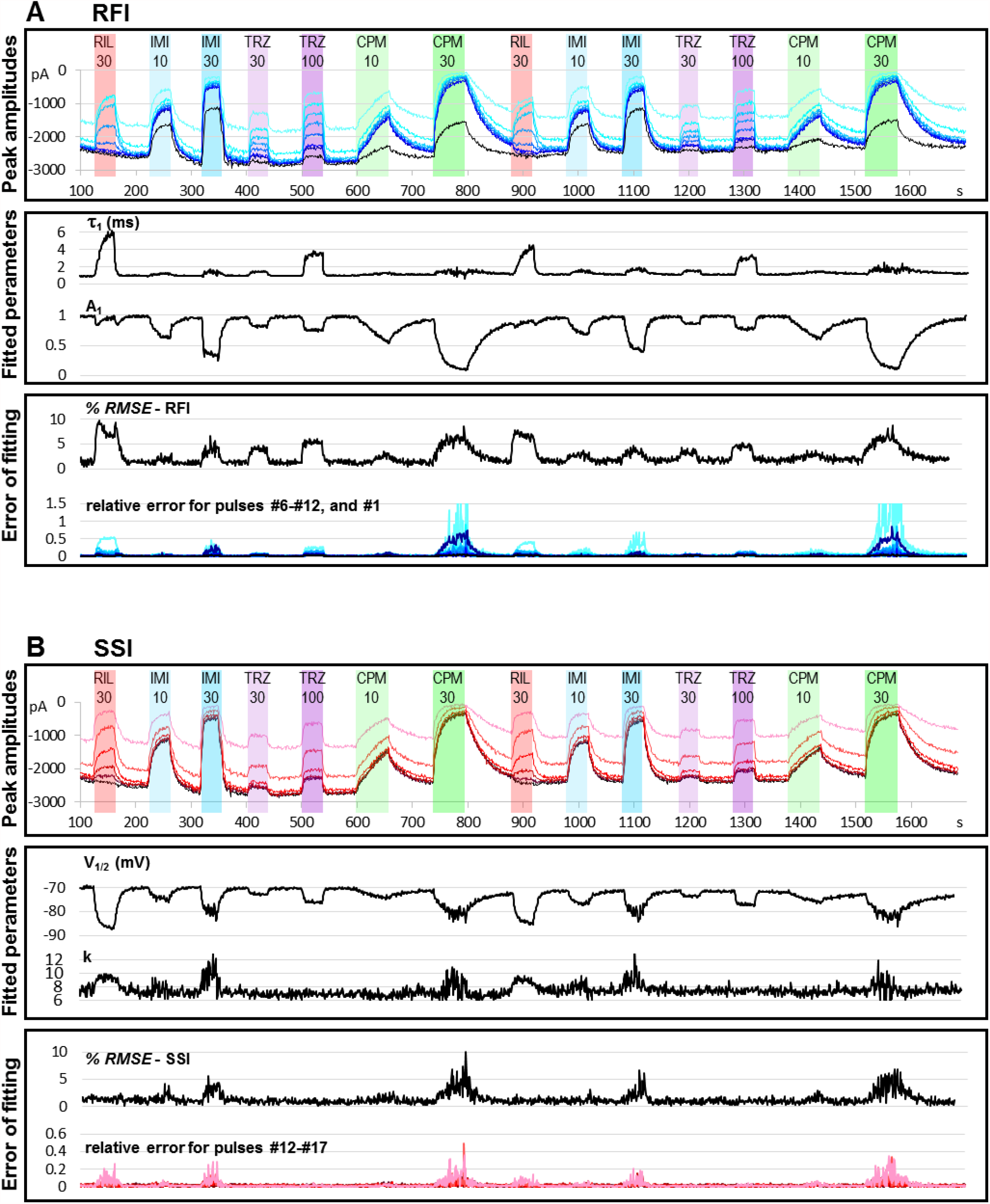
Assessing the accuracy of automated fitting. Peak amplitude plots and fitted parameter plots in both panels A and B are identical with the data shown in Fig. 2 of the paper. Here, however, we also show *RMSE* and *E*_*rel*_ values throughout the whole experiment. Large *E*_*rel*_ values at pulses #7 and #17 are natural, because the fitting procedure minimizes absolute, not relative, squared errors, and these pulses evoke the lowest amplitude currents especially in the presence of an inhibitor. In the case of chlorpromazine and imipramine, we also see a larger relative error for pulse #6. This occurs because although the inhibition after 64 ms hyperpolarization (pulse #6) should be less than the inhibition after 32 ms hyperpolarization (pulse #12), with some of the compounds (imipramine and chlorpromazine) this was not the case (see also Fig. 3D). These compounds have slow micro-dynamics, and therefore they do not only “remember” the 64 ms hyperpolarization, but also that it was preceded by a series of prolonged depolarizations. Riluzole and trazodone, on the other hand, have fast micro-dynamics and have no “memory” of what happened several tens of milliseconds earlier.

**Fig S2.**
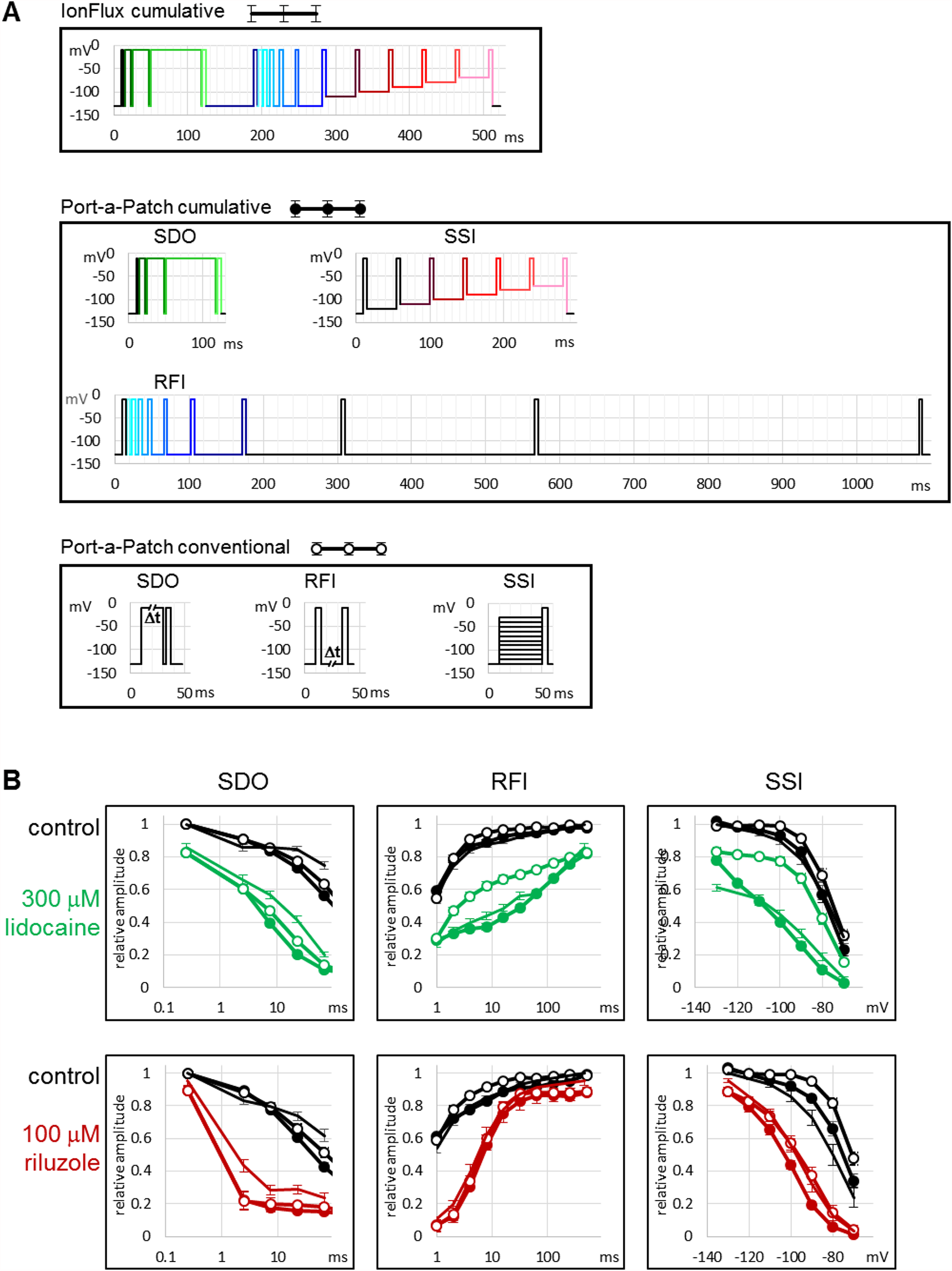
Comparison with single-cell electrophysiology. Port-a-Patch experiments were used to assess the quality of measurements in the IonFlux Mercury instrument, and to compare conventional and cumulative protocols. We sought to answer the following questions: Are the results obtained by the lower seal resistance ensemble recording reliably reflect the behavior of channels measured in single-cell gigaseal recording? Are the cumulative protocols as informative, and as sensitive as conventional protocols? Do the results differ, depending on whether we use the conventional or the cumulative protocols? Three different experimental conditions were compared: Ensemble recordings done by the IonFlux Mercury, using cumulative protocols; single-cell gigaseal recordings done by the Port-a-Patch instrument, using cumulative protocols, and single-cell gigaseal recordings done by the Port-a-Patch, using conventional protocols. The protocols are shown in Fig. S1A Fig. S1B shows the results. Under control conditions there were only minor differences, the SSI curves were somewhat left-shifted in some of the IonFlux experiments. This may be simply due to the fact that these data were recorded in cell ensembles. Ensemble recording measures the sum of 20 parallelly connected cells, therefore there is a chance that not all cells are perfectly sealed. Imperfectly sealed cells may cause the overall voltage control to be less than perfect, or may themselves show left-shifted availability curves, which is a sensitive marker of cell stress. This did not compromise the assessment of drug effects on the whole cell ensemble. In the case of riluzole, where micro-dynamics was fast, IonFlux and Port-a-Patch experiments gave similar results, and there was minimal difference between cumulative and conventional protocols measured with Port-a-Patch. This indicates that micro-dynamics was fast, 40 ms in the SSI protocol was amply enough to reach equilibrium. It was not so, however, in the case of lidocaine, where RFI and SSI plots obtained using conventional vs. cumulative protocols clearly diverged. The 40 ms interpulse interval in the case of the SSI protocols was not enough for reaching equilibrium, in the conventional protocol it was too short for the modulation to develop, while in the cumulative protocol it was too short for recovery. Although this specific protocol is admittedly a result of compromise for the sake of high temporal resolution, we believe that a series of depolarizations occurring at 22.2 Hz from membrane potentials that are close to the resting membrane potential of excitable cells is more relevant with regard to physiological effect than a protocol with prolonged <-100 mV hyperpolarizations. Similarly, in the case of RFI protocols, the cumulative effect of repeated short depolarizations was larger than that of single depolarizations in the conventional protocol, but this increased sensitivity of the protocol may be more relevant in determining the physiological effects of drugs. We can conclude that in the case of drugs with fast micro-dynamics the cumulative protocols are just as effective in assessing properties of inhibition as the conventional ones. In the case of drugs with slower micro-dynamics, we do not believe that cumulative protocols overestimate their potency, but rather that conventional protocols may easily underestimate them if the time periods provided for equilibration are non-physiologically long. Nevertheless, there are compounds where equilibrium cannot be reached even within the 498 ms hyperpolarization between two sweeps. For such compounds lower frequency of testing must be used.

